# The Neurobiology of Cognitive Fatigue and Its Influence on Effort-Based Choice

**DOI:** 10.1101/2024.07.15.603598

**Authors:** Grace Steward, Vikram S. Chib

**Author notes:** Correspondence and requests for materials should be addressed to: Vikram S. Chib, 716 North Broadway, Rm 241, Baltimore, MD 21205, USA.

## Abstract

Feelings of cognitive fatigue emerge through repeated mental exertion and are ubiquitous in our daily lives. However, there is a limited understanding of the neurobiological mechanisms underlying the influence of cognitive fatigue on decisions to exert. We use functional magnetic resonance imaging to examine brain activity while participants make choices to exert effort for reward, before and after bouts of fatiguing cognitive exertion. We found that when participants became cognitively fatigued, they were more likely to choose to forgo higher levels of reward that required more effort. We describe a mechanism by which signals related to cognitive exertion in dlPFC influence effort value computations, instantiated by the insula, thereby influencing an individual’s decisions to exert while fatigued. Our results suggest that cognitive fatigue plays a critical role in decisions to exert effort and provides a mechanistic link through which information about cognitive state shapes effort-based choice.

## INTRODUCTION

The workday is filled with tasks that require a great deal of cognitive effort, whether that be meeting with clients and coworkers, preparing for presentations, or simply replying to emails. As the day progresses, our engagement in these effortful tasks leads to fatigue, which impacts our choices by making us less willing to exert cognitive effort. Recent studies in cognitive neuroscience have begun to dissect the circuitry underpinning physical fatigue and its influence on effort-based decision-making^1,2^. While previous works have examined the effects of cognitive fatigue on task engagement and performance^3,4^, there is a limited understanding of how the emergence of fatigue during repeated cognitive exertion influences effort-based decision-making at the level of brain and behavior.

Experiments examining the influence of physical fatigue on effort-based decision-making have shown that fatigued individuals increase their subjective costs of physical effort, and higher rewards are needed to incentivize exertion while in a fatigued state^1,2,5^. Physical effort costs and effort decisions fluctuate with individuals’ momentary fatigue^5,6^. Neuroimaging during exertion and choice has revealed a network of brain activity that underlies effort-based choices and is impacted by physical fatigue^1,2,7–9^. This network includes the anterior cingulate cortex (ACC), bilateral anterior insula, and ventromedial prefrontal cortex (vmPFC), which are integral for computing the value of effortful options and making effort-based decisions. It has also been shown that during physical effort-based decision-making, the right anterior insula (rIns) is sensitive to fatigue-induced changes in prospective effort and is related to exertion signals in the premotor cortex^1^. A study that examined physical exertions directly interleaved with choice also showed that the medial and lateral prefrontal cortex tracked individuals’ states of fatigue and that a frontostriatal network integrated fatigue with effort value^2^.

One important factor in signaling fatigue is the ability to sense one’s internal state. Notably, the rIns is responsible for encoding proprioceptive signals, and the anterior insula encodes effort value^1,10–12^. We recently showed, that in the context of physical effort-based decision-making, feelings of fatigue mediate effort values encoded by the insula^1^. However, previous studies that examined neural signals at the time of exertion and rest were focused on physical effort and were not designed to examine prospective cognitive effort valuation and fatigue^2–4,13,14^. Furthermore, while recent studies have examined how physical and cognitive fatigue influence cognitive control during a temporal discounting task, they did not evaluate how fatigue influences cognitive effort valuation^15,16^. Therefore, it is unclear how brain signals related to cognitive fatigue might influence effort valuation and whether similar functions of rIns are involved in signaling cognitive fatigue and cognitive effort-based choice. To our knowledge, there have been no studies that have directly tested how bouts of cognitive exertion influence brain signals related to the prospective valuation of cognitive effort and resulting decisions. As such, there is a limited neurobiological understanding of how cognitive fatigue influences effort valuation and decisions to engage in cognitive exertion.

In this study, we investigated the influence of cognitive fatigue on behavioral representations of subjective effort value and the neural mechanisms by which fatigue influences the brain’s decision-making circuitry. We hypothesize that fatigue, resulting from repeated cognitive exertion, will reduce individuals’ willingness to exert. In a fatigued state, compared to a rested state, when individuals face the option of exerting greater prospective cognitive effort for reward, they will be less willing to accept high-effort options. This hypothesis has its basis in previous studies of physical effort-based decision-making that found that individuals exhibit increased costs of physical effort when in a fatigued state^1,2^. We hypothesize that decisions about prospective cognitive effort exertion have their basis in a value signal encoded in the ACC and insula. This hypothesis is informed by neuroimaging studies, which consistently show that activity in regions of the ACC and insula is related to subjective effort value^1,7,10,17,18^. Given our recent study of physical fatigue, which showed that rIns was sensitive to fatigue-induced changes in effort-based decision-making^1^, we predict that the insula is also sensitive to changes in cognitive effort value as a function of cognitive fatigue. We predict that brain regions related to cognitive exertion will be functionally coupled to rIns during choice, suggesting a network through which exertion and fatigue are translated into subsequent effort choices. These hypotheses form a neurobiological framework of cognitive fatigue, which recruits brain regions responsible for effort valuation and cognitive exertion to inform decisions about prospective effort while fatigued.

## RESULTS

There are many different cognitive tasks for which repeated engagement induces feelings of fatigue. In this study, we chose to focus on cognitive effort in the form of an *n*-back working memory task because it allows for clearly operationalized levels of exertion and well-mapped engagement of task-induced neural activity^17,19^. During the working memory task, participants attended to a serial presentation of letters, and attempted to identify when the letter presented matched the one *n* letters previously presented (Figure 1A). Additionally, each effort level was assigned a unique color so that during choice trials, the cognitive effort level could be cued without explicitly presenting the numerical *n*-back level. This was meant to minimize the association of cognitive effort with the *n*-back number and instead increase reliance on the participants’ subjective feelings of cognitive effort.

**Figure 1.**
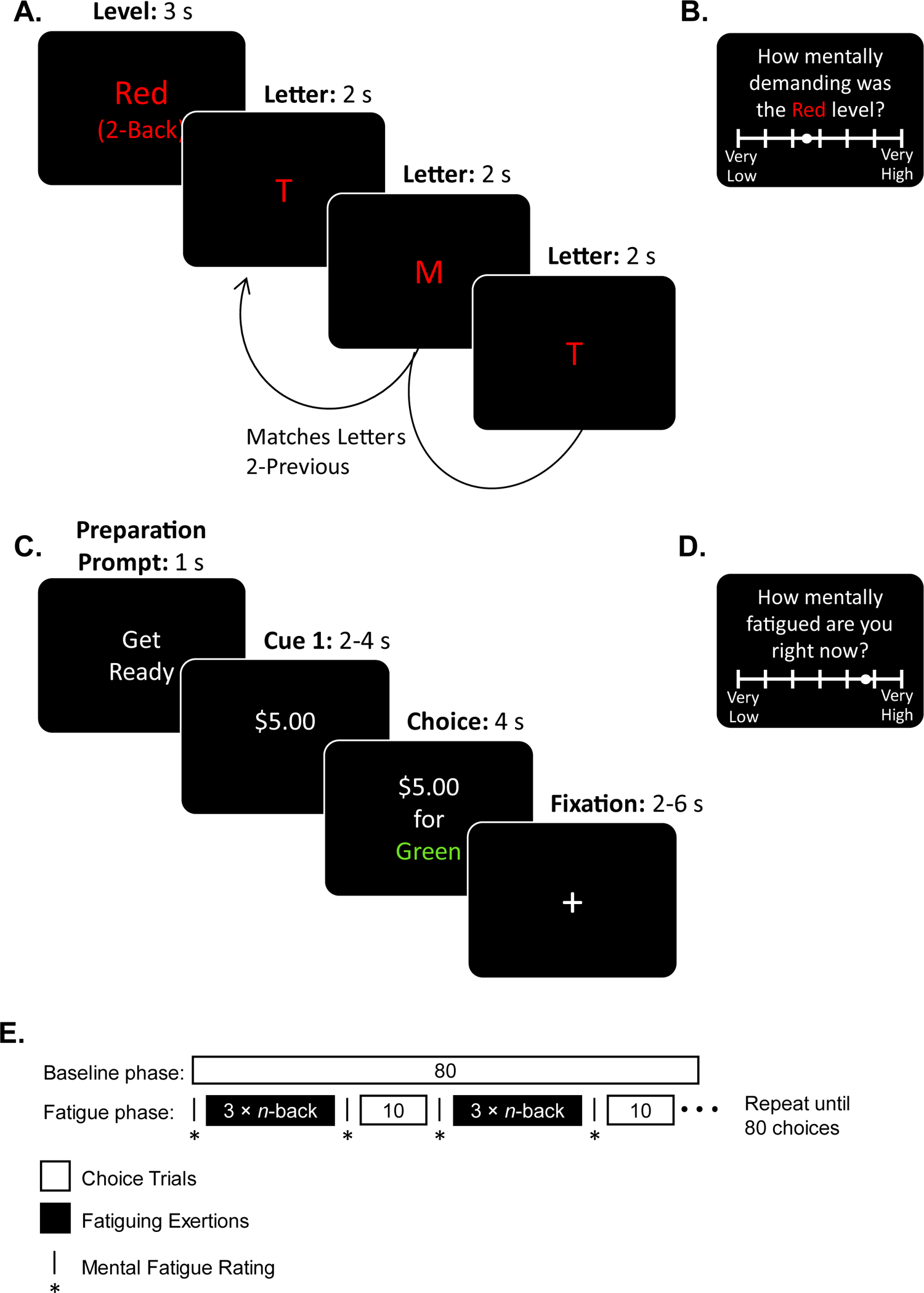
Experimental Design. **A.** Participants first associated different effort levels of *n*-back task with distinct colors. During the *n*-back cognitive effort task, participants identified when a letter presented on the screen matched the one presented *n* letters previous. **B.** Following this association phase, participants rated the difficulty of each color level on a continuous scale from ‘Very Low’ to ‘Very High’ mental demand. **C.** During choice trials, participants made decisions between a default effort/reward (1-back for $1) option or an option with varying levels of effort for different reward amounts. Each trial began with the presentation of one piece of information regarding the non-default option (either effort level or reward amount), followed by presentation of the second piece of information. Participants were instructed to make their choice after the second piece of information was presented. **D.** Participants rated their state of mental fatigue on a continuous scale from ‘Very Low’ to ‘Very High.’ This rating was repeated several times during the fatigue phase. **E.** Experiment Schedule. To study the effects of fatigue on effort-based decision-making, blocks of cognitive exertion trials were interspersed with blocks of effort-based choice trials. The experiment was divided into baseline and fatigue choice phases, which were both scanned with fMRI. The baseline choice phase consisted of 80 effort-based choices presented in a pseudo-randomized order. Following the baseline choice phase, participants performed the fatigue choice phase of the experiment, in which they underwent repeated exertion trials (indicated in black) to bring them into a fatigued state. The fatigue choice phase was comprised of alternating cognitive exertion and choice blocks. Cognitive exertion trials involved performing a 3-back working memory task three times. Each choice block consisted of 10 effort-based choices randomly sampled from the same set used in the baseline choice phase. At the beginning and end of each exertion block participants rated their cognitive fatigue. Completion of the fatigue choice phase consisted of 8 back-to-back exertion and choice blocks.

To develop an association between cognitive effort levels and color cues, participants were randomly presented with *n*-back tasks from levels 1 through 6, with their associated colors (Figure 1A). To ensure that participants had developed an association between the extent of cognitive effort associated with each color cue, participants rated the mental challenge of each *n*-back level based on the color cue alone, following the assessment trials (Figure 1B).

To study how decisions about prospective cognitive effort are influenced by cognitive fatigue, we scanned participants’ brains with functional magnetic resonance imaging (fMRI) while they made choices to exert different levels of cognitive effort for varying amounts of money, interspersed with bouts of fatiguing cognitive exertion (Figure 1C). First, during a baseline choice phase, participants made choices to either accept a default option of a 1-back task for the reward of $1, or to accept a variable higher-level task for a greater reward (non-default option). These choices were meant to characterize a person’s decision-making before fatigue. This baseline phase was immediately followed by a fatigue choice phase, during which participants completed alternating blocks of fatiguing working memory exertions and the same effort-based choices. Participants rated their mental fatigue at the beginning and end of each exertion block (Figure 1D). All choices were for prospective effort and reward, and at the end of the experiment, two choices were randomly selected from the baseline and fatigue choice phases and played out.

### Fatigue-induced changes in effort-based choice

To confirm that participants understood the association between cognitive effort levels and color cues, we examined the relationship between participants’ ratings of mental demand and the *n*-back-associated color cues. We found that participants mental demand ratings increased with cues associated with higher *n*-back levels (linear mixed effects model; t_166_ = 12.78, p = 1.91E-26, Figure 2A, Table S1). This correlation between ratings of mental demand and cognitive effort levels indicates that participants understood the mapping between feelings of cognitive exertion and effort color cues. As a result, participants could make meaningful choices about prospective effort options that utilized these color cues during the choice phases.

**Figure 2.**
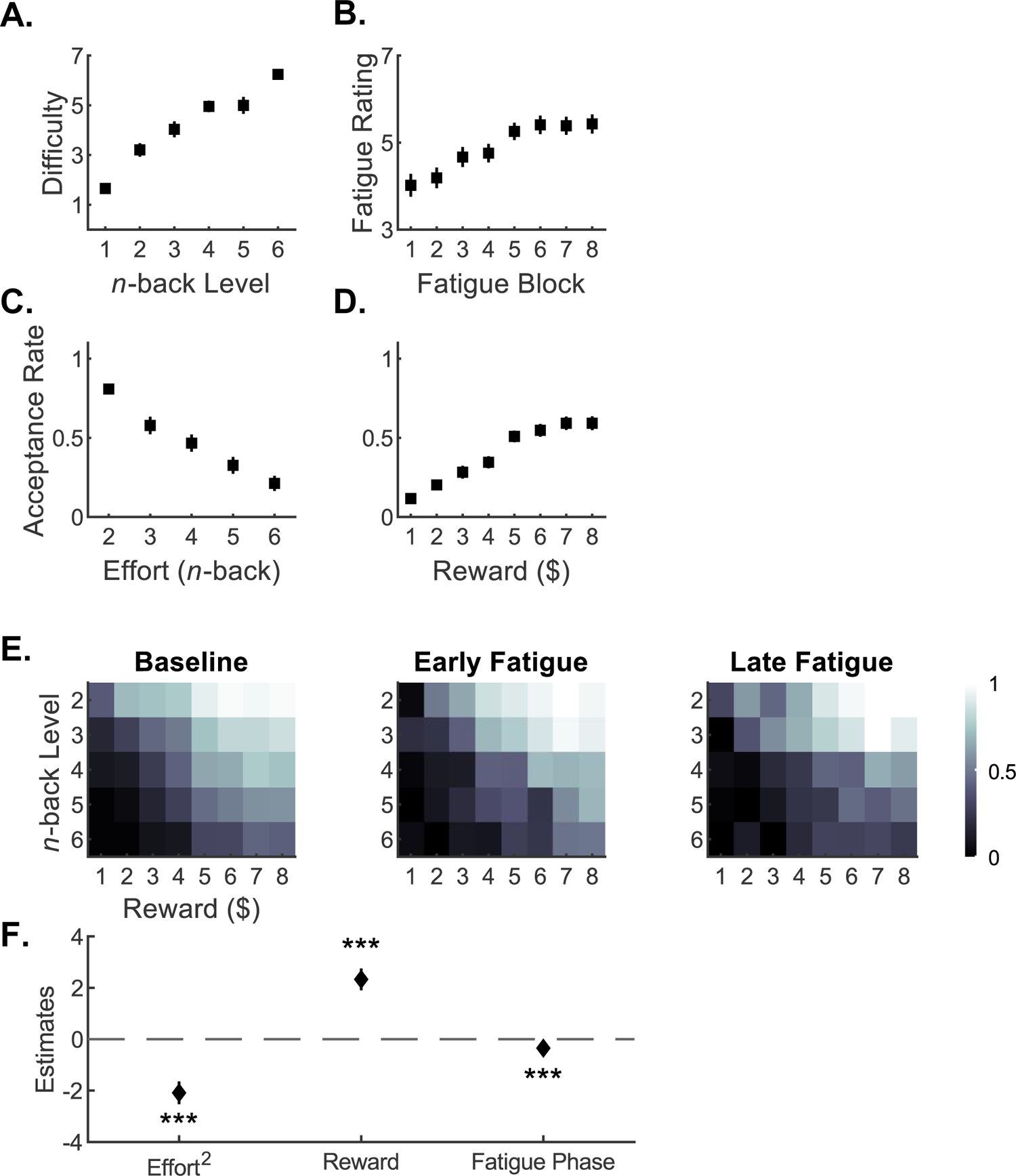
Effects of Fatigue on Choice. **A.** Participants ratings of working memory task difficulty as a function of the *n*-back level, during trials polling mental demand. **B**. Ratings of fatigue during the fatigue phase, as a function of exertion block. Probability of accepting the non-default effort/reward option as a function of cognitive effort level (**C**) and reward (**D**). Error bars denote SEM. **E.** Proportion of non-default option choices accepted, across all participants, as function of effort level and reward at different time points in the experiment. Participants were more likely to choose options with higher reward and lower effort. The main difference in behavior between the baseline and fatigue phases was at the highest levels of offered reward and effort. **F.** To better understand the effect of fatigue on behavior, we used a generalized linear mixed effects model and found a significant effect of the baseline/fatigue condition on choice. When participants became fatigued, they were more likely to take the default option and forgo greater amounts of reward for increased effort. Error bars represent SE. *** p < 0.001.

To verify that participants became cognitively fatigued through repeated exertion during the fatigue phase, we examined how their mental fatigue ratings changed as a function of cognitive exertion blocks. We found that as participants progressed through the fatigue blocks, their rating of mental fatigue increased (linear mixed effects model, t_222_ = 6.95, p = 3.94E-11, Figure 2B, Table S2). We also examined the relationship between participants’ performance and exertion block. We did not find evidence that participants’ task performance declined with additional fatigue. Instead, there was a significant increase in performance with additional fatigue blocks (linear mixed effects model; t_222_ = 6.95, p = 3.90E-11, Figure S1, Table S3). These results illustrate that repeated cognitive exertion of the working memory task leads to increased feelings of fatigue without resulting in decrements in performance.

Next, we analyzed choice trials across both the baseline and fatigue phases. We found that individuals’ willingness to accept the non-default effort/reward option decreased as cognitive effort levels increased (linear mixed effects model; t_138_ = −9.18, p = 5.65E-16, Figure 2C, Table S4) and increased with additional reward (linear mixed effects model; t_222_ = 8.96, p = 1.36E-16, Figure 2D, Table S5). These results align with several previous studies of cognitive and physical effort-based decision-making, which found that increasing effort and decreasing reward have inverse effects on effort-based choice^5,10,17,20,21^.

To evaluate the effect of fatigue on effort-based choice, we analyzed the difference between individuals’ propensity to accept the non-default effort option in the baseline and fatigue phases. We found that individuals tended to accept fewer non-default option trials in the fatigue phase compared to baseline and that this propensity appeared to increase later in the fatigue phase (Figure 2E). As in previous studies of effort-based decision-making^10^, we used a parameter estimation procedure to fit models of subjective value for each participant and found that a parabolic effort discounting function best described participants’ choice (Figure S2). To formally test the effect of fatigue on effort-based choice, we examined the relationship between participants’ propensity to accept the non-default subjective effort/reward options as a function of effort and reward magnitude and experimental phase (baseline/fatigue). We found a decreased acceptance of the non-default effort/reward option in the fatigue phase relative to baseline (generalized linear mixed effects model; β_Fatigue_ = −0.349, SE = 0.097, t_4434_ = −3.60, p = 3.24E-4, Figure 2F, Table S6) in a model that performed better than the null model with no effect of phase (LR = 12.773 > χ^2^(1) = 3.841). When fatigued, participants preferred the low-effort, low-reward default option compared to when in a rested state.

### Neural encoding of cognitive effort value

We found that the BOLD signal in a network of brain regions, including the dorsal anterior cingulate cortex (dACC), nucleus accumbens (NAc), ventromedial prefrontal cortex (vmPFC), and right anterior insula (rIns), was modulated by the subjective chosen value of the effort/reward option across both the baseline and fatigue phases (Figures 3A, 3B). This finding is consistent with several studies of effort-based decision-making for cognitive and physical effort that implicated these brain regions as part of an effort valuation network^10,17,18^.

**Figure 3.**
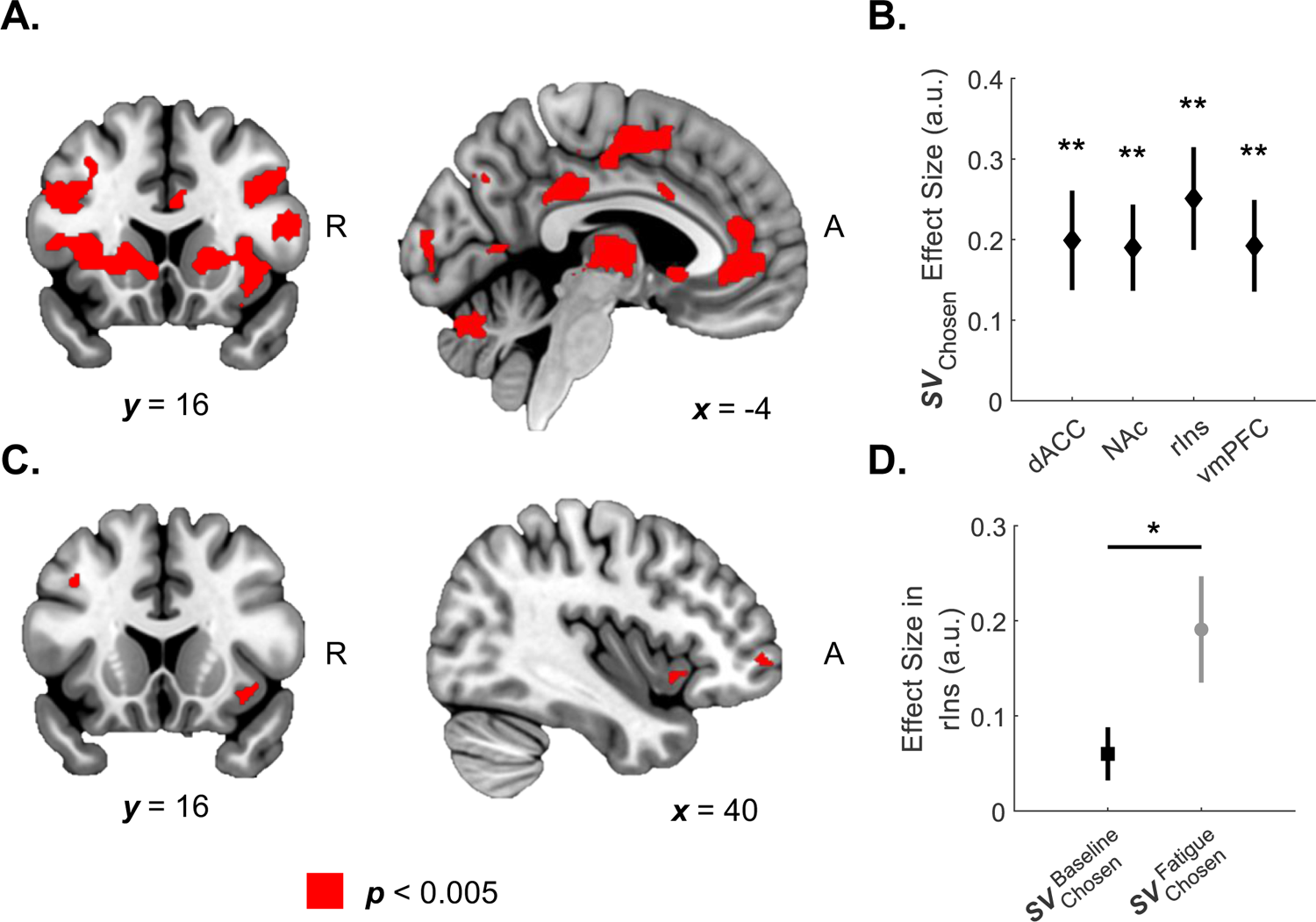
Neural Representations of Chosen Value. **A**. Overall chosen value encoding. Whole brain activity thresholded at voxelwise p < 0.005 with an extent threshold of 10 voxels. **B.** Activity in the dorsal anterior cingulate cortex (dACC: peak = [−8, −2, 54], t27 = 3.22, p = 0.007), nucleus accumbens (NAc: peak = [18, 8, −2], t27 = 3.55, p = 0.003), right anterior insula (rIns: peak = [36, 26, −12], t27 = 3.94, p = 0.001,), and ventral medial prefrontal cortex (vmPFC: peak = [−8, 42, −2], t27 = 3.38, p = 0.004) encoded the chosen subjective value (computed from subjective effort value and reward) across both the baseline and fatigue phases. Effect sizes in a priori regions of interests (ROIs) for chosen subjective value across the baseline and fatigue choice phases are shown and Bonferroni corrected across the 4 ROIs. Error bars indicate SEM. **p < 0.01 **C.** Whole brain analysis contrasting chosen subjective value encoding between fatigue and baseline phases thresholded at voxelwise p < 0.005 with extent threshold of 10 voxels. **D.** Activity in rIns (peak: [34, 16, −12], t27 = −2.14, p = 0.042) reflects the difference in chosen and unchosen subjective value, between the baseline and fatigue phase via ROI analysis in an a priori region of interest. Error bars indicate SEM. *p < 0.05

To test for brain regions sensitive to changes in chosen subjective value induced by cognitive fatigue, we contrasted the difference between chosen subjective values in the baseline and fatigue phases. We found that a region of the rIns had increased sensitivity to the chosen subjective value during the fatigue phase, compared to baseline (Figures 3C, 3D). This suggests that activity in rIns is sensitive to changes in subjective effort value resulting from cognitive fatigue, which aligns with previous studies of effortful exertion that have suggested that this brain region encodes representations of individuals’ internal state as a function of physical exertion and fatigue^1^.

### Cognitive fatigue-induced changes in working memory neural activity

We examined how cognitive fatigue influenced brain activity during the fatiguing working memory task. We found that as fatiguing *n*-back blocks increased, the signal in the bilateral dorsolateral prefrontal cortex (dlPFC) also increased (Figures 4A, 4B, Table S8). Activity in this region has been shown to increase with increasing working memory load (i.e., increasing *n*-back level)^22,23^. With this in mind, increasing blocks of exertion in the context of our fatigue paradigm seem to increase neural representations of cognitive load.

**Figure 4.**
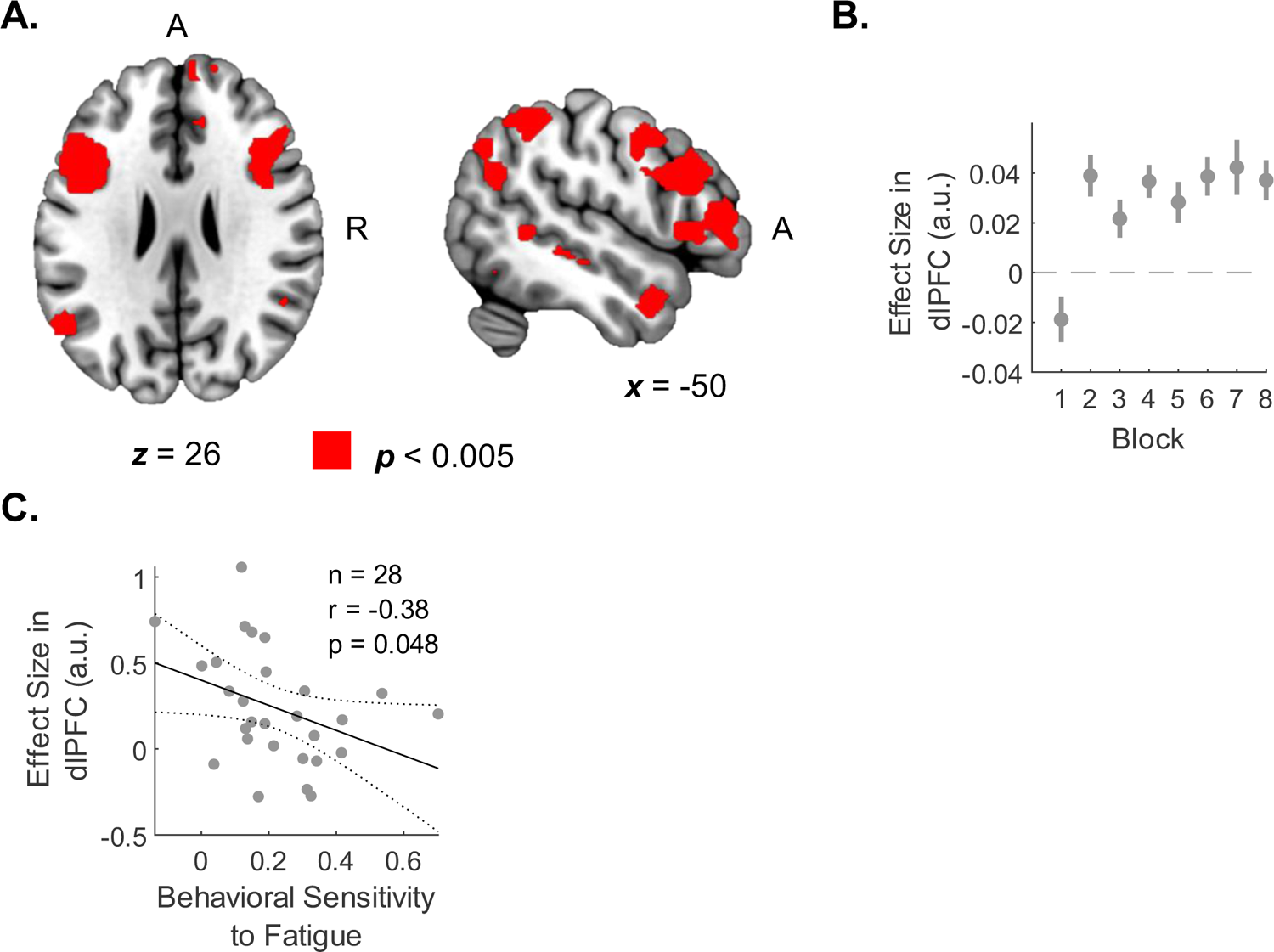
Neural Signatures of Cognitive Exertion Induced Fatigue. **A.** *n*-back exertion induced changes in brain activity. Activity in bilateral dlPFC increased as a function of exertion block, at the time of fatiguing exertions. We estimated a general linear model with a parametric modulator for each fatiguing block completed and found a significant increase in activation after the first fatigue block in areas previously associated with working memory function (L dlPFC peak: [−38, 4, 32], R dlPFC peak: [4, 8, 28], t27 = 3.80, p = 7.5E-4). **B.** Illustration of the effect size in bilateral dlPFC ROI increasing with additional fatiguing blocks. Error bars indicate SEM. **C.** We found a significant correlation between the slope of participants’ fatigue ratings during the fatigue phase (Behavioral Sensitivity to Fatigue) and decreased activation in the bilateral dlPFC ROI.

Considering our prediction that cognitive state would be an integral factor in representations of cognitive fatigue, we tested whether individuals’ ratings of cognitive fatigue were indexed by fatigue-induced changes in activity in the dlPFC. It has been suggested that fatigue might arise due to discrepancies between expectations about cognitive exertion and the actual neural processing required to achieve the exertion^24^. In the context of our experiment, if an individual’s working memory system does not adjust neural activity in response to repeated cognitive exertions, one might feel that effort is particularly costly because of the discrepancy between the cognitive exertion that one believes they can achieve and their actual exertion. In contrast, it is possible that fatigue arises from an accurate representation of an individual’s cognitive state and that changes in feelings of fatigue are simply a reflection of the altered neural activity that occurs following fatiguing exertion. We calculated a measure of participants’ behavioral sensitivity to fatigue as the slope between fatigue block number and their fatigue ratings. Using each participant’s behavioral sensitivity to fatigue as a covariate with activity in dlPFC, we found that those individuals with greater behavioral sensitivity to fatigue exhibited less change in dlPFC activity (Pearson correlation, *r* = −0.38, p = 0.048, Figure 4C). These findings are consistent with the idea that miscalibrated recruitment of neural activity in cognitive regions is related to increased feelings of fatigue — those participants who report the highest changes in fatigue are those who do not modify their dlPFC activity to accommodate the reduced cognitive capacity that results from repeated mental exertion. These results align with our previous findings in the domain of physical fatigue, in which a lack of calibration of motor cortical signals was related to increased feelings of fatigue^1^.

### Functional Connectivity Between rIns and dlPFC

Finally, given our hypothesis that information about one’s cognitive state is integral to assessing cognitive capacity and associated effort value, we tested the idea that the neural circuit modulating cognitive effort value representations in rIns might be influenced by computations about cognitive exertion instantiated in dlPFC during choice. To test this hypothesis, we conducted a psychophysiological interaction (PPI) analysis between rIns (seed) and dlPFC (target) with fatigue state as a psychological variable (Figure 5A). This analysis revealed a robust modulation of connectivity between the dlPFC and rIns as a function of fatigue state at the time of choice (Figure 5B). Connectivity was increased in the fatigued state compared to baseline, and in the fatigued state, activity in the rIns at the time of choice was associated with increased connectivity to the dlPFC. These results provide support for the hypothesis that activity in dlPFC and rIns are functionally related during effort-based decision-making and suggest that interactions between these brain regions could facilitate the transfer of information about cognitive state that is used to inform choices about prospective cognitive effort when cognitively fatigued.

**Figure 5.**
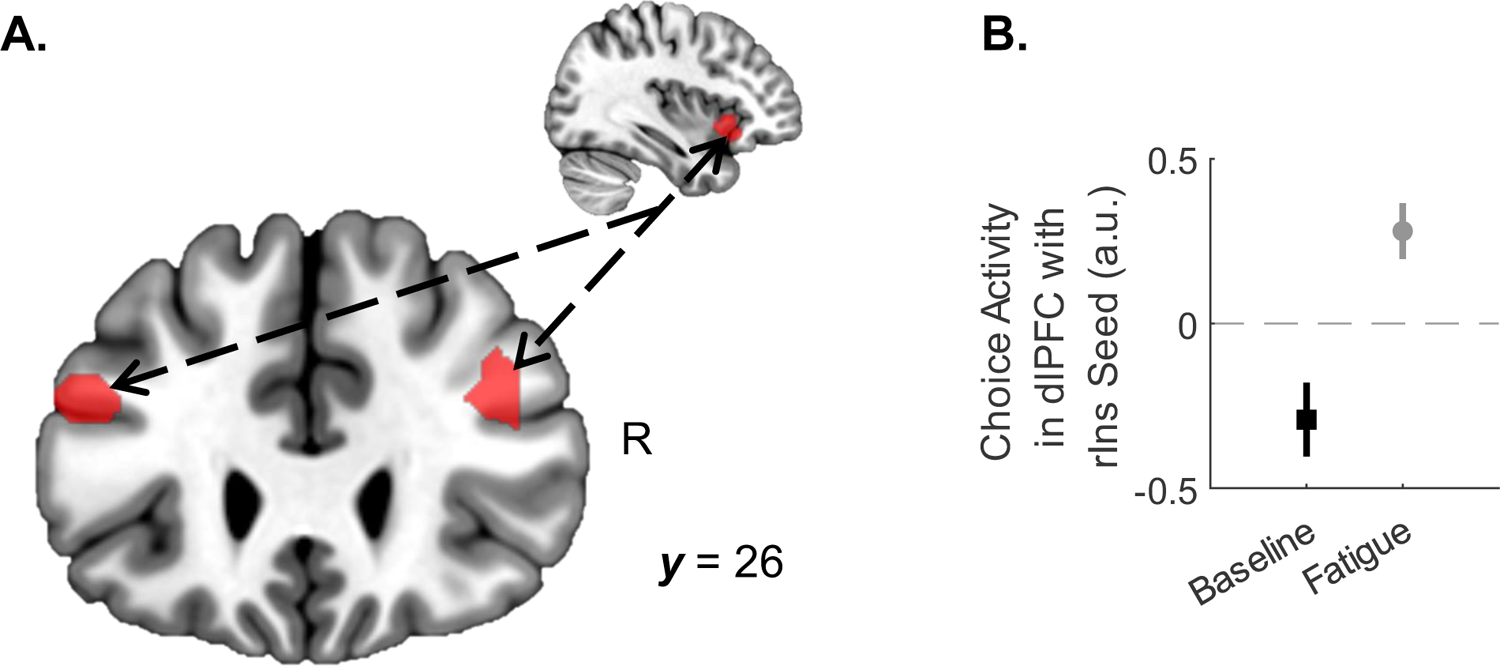
Functional Connectivity between rIns and dlPFC. **A.** Illustration of the psychophysiological interaction (PPI) analysis. We computed a PPI between rIns and bilateral dlPFC with the psychological variable of baseline/fatigue state, at the time of choice. Activity in bilateral dlPFC shows an increased functional coupling with rIns during the Fatigue Choice Phase, compared to the Baseline Choice Phase (L dlPFC peak = [−44, 0, 30], R dlPFC peak = [34, −2, 42], t27 = 3.92, p = 5.52E-4). Statistical inference was determined via an ROI analysis of the contrast of the two phases. **B.** Effect size plots by phase of rIns dlPFC functional coupling. Effect size plots for each phase are for visualization purposes only. Error bars denote SEM.

## DISCUSSION

In this study, we show that cognitive fatigue decreases an individual’s willingness to exert higher cognitive effort, that decisions to exert effort while fatigued are related to value signals in the rIns, and that signals related to cognitive exertions in dlPFC are functionally coupled with signals in rIns at the time of choice. Our neuroimaging findings are consistent with previous studies showing that effort value signals are represented in a network of brain regions, including the rIns, and that activity in rIns follows the time course of feelings associated with exertion and rest ^1,17^. However, previous studies primarily focused on physical effort and fatigue and did not examine how cognitive fatigue is related to decisions to exert effort. Our results build on these previous studies by illustrating a mechanism by which signals related to cognitive fatigue-modulated effort value are shaped by brain regions responsible for generating fatiguing cognitive exertion.

Previous studies of effort cost have mainly focused on the trade-off between prospective physical effort (e.g., grip exertion, button pressing) and reward ^1,2,7,18^ and have implicated a network of brain regions, including the vmPFC, ACC, and insular cortex. Studies examining trade-offs between cognitive effort (e.g., working memory tasks, arithmetic) and reward have identified a network that largely overlaps with physical effort, suggesting a general effort valuation network that subserves decisions to exert^10,17,25^. Our results are consistent with these findings, which identify regions in the ACC and rIns responsible for effort-reward trade-offs in the context of cognitive effort decisions in both rested and fatigued states. Neuroimaging studies of the influence of fatigue on decisions to exert have focused on the domain of physical effort and shown that regions of the prefrontal cortex, rIns, and motor cortex, are sensitive to fatigue state and influence decisions about effort exertion. Here, we show that a region of rIns previously shown to be sensitive to physical chosen effort value when physically fatigued is also sensitive to cognitive chosen effort value when cognitively fatigued^1^. Moreover, these regions are functionally coupled with working memory cognitive exertion-related signals in dlPFC at the time of choice, which suggests that information about cognitive state could influence these choices. These results point to a general role of rIns in being sensitive to feelings of fatigue while experiencing both cognitive and physical fatigue.

We found that dlPFC signals related to fatiguing exertion were inversely related to the extent to which participants rated fatigue – those individuals who became more fatigued through repeated exertion exhibited smaller changes in fatigue-induced dlPFC activity. These results are consistent with the idea that cognitive fatigue may be associated with dyshomeostatic representations of cognitive exertion in dlPFC^1^. Those individuals who do not tune their dlPFC activity following cognitively fatiguing exertion find prospective effort to be more costly. The cognitive effort might feel particularly costly to these individuals because they continue to recruit the same levels of neural activity as in a baseline state, although they are physiologically unable to recruit higher levels of neural resources to efficiently achieve the target levels of cognitive performance. This idea aligns with recent theoretical accounts of fatigue that suggest discrepancies between perceptions of ability and actual sensorimotor capacity may give rise to feelings of fatigue^26^.

Recent studies have shown that neurotransmitters such as gamma-aminobutyric acid (GABA) and glutamate play a key role in judgments of physical effort and are related to cognitively fatiguing exertion^27,28^. For physical effort, GABA inhibition has been related to changes in descending motor drive and variability in motor output, neural and behavioral signatures that impact feelings of exertion^27^. However, the neurochemical underpinnings of cognitive effort are less straightforward. Working memory tasks that result in greater cognitive fatigue and require greater cognitive control have been associated with higher glutamate concentration and glutamate/glutamine diffusion in task-related regions^29^. It has been proposed that these neurochemicals could be associated with recycling potentially toxic substances accumulated in the brain during sustained cognitive exertion. The neural system’s ability to monitor these state changes could be integral in signaling and updating feelings of fatigue.

In conclusion, our study provides evidence of how the brain integrates information about cognitive fatigue into effort valuation and decisions to exert. Our data implicate a mechanism by which cognitive exertion-related signals in dlPFC influence effort valuation signals in the rIns, which underlie decisions to exert while in a fatigued state. The present work begins to bridge the gap in understanding how the brain represents cognitive fatigue and the influence of cognitive fatigue on decisions to exert, which to date has received limited study. Moreover, from a clinical standpoint, the neurobiology of cognitive fatigue is likely to prove important in understanding feelings of amotivation that are prevalent across neurological and psychiatric conditions. From this perspective, the behavioral and neural underpinnings of fatigue provide a foundation for the development of interventions aimed at optimizing effortful exertion.

## ACKNOWLEDGEMENTS

This work was supported by the Eunice Kennedy Shriver National Institute of Child Health & Human Development of the National Institutes of Health under Award Number R01MH119086 and the National Institutes of Mental Health R01HD097619.

## METHODS

### Participants

The Johns Hopkins School of Medicine Institutional Review Board approved this study. Participants were right-handed, had no self-reported history of psychiatric or neurological disorders, and had no self-reported previous adverse experiences undergoing an MRI exam. All participants provided informed consent before beginning the study.

Thirty-four individuals participated in this study. Two participants were excluded for poor performance on the *n*-back task (not identifying at least 50% of targets on average), two were excluded for making choices with no variability (i.e. accepting all offers), and two were excluded for having ratings of *n*-back difficulty that declined with increasing *n* (i.e. more difficult *n*-back tasks were not perceived as more demanding). This left a final cohort of n = 28 participants (mean age 25 years; standard deviation in age, 4 years; 18 females).

### Paradigm

Before the experiment, participants were informed that they would receive a $50 show-up fee and that the opportunity for additional monetary earnings was dependent on their choices and task performance. This customized experimental procedure was built in MATLAB 2020b^30^ using Psychophysics Toolbox Version 3^31^ for stimuli presentation and behavioral data collection.

The experiment began with an association phase in which participants performed different cognitive effort levels of an *n*-back working memory task (Figure 1A). In this paradigm, cognitive effort was operationalized as the level of the *n*-back task — higher *n* corresponded to higher cognitive effort. Each *n*-back trial was comprised of 40 sequentially presented letters. Within this sequence were 10 target letters that a participant would have to identify as the same letter *n* letters prior. Participants had 2 seconds to identify whether the current letter on the screen was one of these targets. Once the participant entered their choice, the next letter was presented. Participants had to correctly identify 50% of the targets to succeed in the task. Six levels of *n*-back were presented (n = 1-6), a distinct color was pseudo-randomly assigned to each level, and the stream of sequential letters was presented in this color. Participants were presented with varying levels of *n*-back in blocks of three trials. Each level was presented in random order without replacement.

Following this association phase, participants were randomly presented with color cues that had previously been associated with *n*-back levels and were instructed to rate their perceived level of mental demand (Figure 1B). Ratings were made on a continuous scale from “Very Low” to “Very High”.

To examine the effect of cognitive fatigue on behavioral and neural representations of effort valuation, we scanned participants’ brains with fMRI while they made decisions about prospective cognitive effort and monetary reward, before and after they performed repetitive fatiguing cognitive exertions. Before being presented with the effort/reward decisions, participants were told that two of their decisions would be randomly selected (one from decisions before and one from after cognitive exertions) and played out at the end of the experiment. Since trials were extracted at random, participants were instructed that they did not need to spread their exertions over all their trials and that they should treat each effort decision individually.

During the baseline choice phase, which was meant to elicit effort and reward preferences in a rested state, participants were presented with a series of effort/reward choices between performing a 1-back task for $1 (default option) or a higher level of cognitive effort for reward (non-default option). Participants made their choices by pressing one of two buttons on a handheld button box with their right hand (Cedrus Corporation, San Pedro, CA, Cedrus RB-830). For the non-default options, the effort levels were drawn from the six color cues introduced in the association phase, and rewards ranged from $1 to $8 in $1 increments. Effort/reward options were presented consecutively in a pseudo-random order, such that choices of approximately equal choice difficulty were shown in every block of 10 choices. This phase consisted of 40 unique choices. Offers were presented to participants twice: once with the non-default effort presented first and once with the reward presented first. This allowed for the possibility of analyzing the separate valuation of effort and reward before the time of choice. Offers were presented in a pseudo-random order, such that choices of approximately equal difficulty were shown in every block of 10 choices to mitigate offer order presentation as a confound.

A fatigue phase immediately followed the baseline phase. During this fatigue phase, participants were asked to rate their mental fatigue on an identical continuous scale from “Very Low” to “Very High” with 7 tick marks on the number line to act as a reference. Participants then underwent a fatiguing trial, where they performed three 3-back tasks. They rated their mental fatigue again before going on to make 10 choices. This process was repeated a total of 8 times, such that identical offers were given to the participant during both phases. Participants were made aware that they had to succeed on 50% of all fatiguing trials to receive the opportunity to earn additional rewards at the end of the experiment during the outcome phase.

Following imaging, the participant choices were realized. If the participant had met the fatigue phase success threshold, two offers from the baseline and fatigue phases were chosen randomly. The participant then completed their chosen option, exerting the effort required and receiving the associated reward. Participants had to achieve the effort task to receive the reward and were given three opportunities to succeed.

### MRI Protocol

We conducted our MRI scans using a Philips dStream Achieva 3T TX scanner equipped with a DirectDigital RF system. Participants lay supine on the MR exam table and had their head stabilized within a 32-channel SENSE head coil, whose signal was amplified by an 18 kW Solid-state RF power amplifier.

High-resolution structural images were collected using an MPRAGE T1 weighted pulse sequence with an Echo Time of 3.57ms and Repetition Time (TR) of 8.12ms. The resulting whole-brain images had a 1mm × 1mm × 1mm voxel resolution.

Functional images were collected at an angle of 30° from the anterior commissure–posterior commissure (AC–PC) axis, which reduced signal dropout in the orbitofrontal cortex^32^. Functional T2 weighted images included 48 slices in 1.875mm x 1.875mm x 3mm resolution. We used an echo-planar imaging (FE EPI) pulse sequence (TR = 2800 ms, TE = 30 ms, FOV = 240, flip angle = 70°).

### Mixed-Effects Models of Behavior

To evaluate participants’ ability to recall the cognitive effort associated with each color level. we employed a linear mixed-effects model:

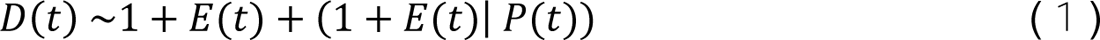

Here, *D(t)* denotes the cognitive difficulty of the assessed effort level, *E(t)* represents the *n* in the *n*-back being rated, and *P(t)* is a categorical participant identifier for a given trial *t*. Both *D(t)* and *E(t*) were z-scored using the *zscore* function in MATLAB 2023a^33^. Models were fitted via maximum likelihood estimation using the *fitlme* function with default settings, incorporating a random effect of the slope to account for varying participant sensitivity to increasing levels of the *n*-back task.

To evaluate how participants’ fatigue ratings increased with additional fatiguing blocks, using the following model:

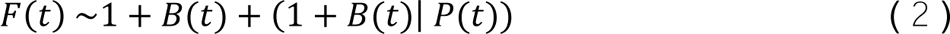

In this context, *F(t)* represents participants’ ratings of mental fatigue after completing exertion block *t*, *B(t)* denotes the cumulative number of blocks completed, and *P(t)* is a categorical participant identifier. *F(t)* and *B(t)* were both z-scored. Models were fitted via maximum likelihood estimation, introducing a random effect of the slope of the fatigue rating to accommodate varying participant sensitivity to additional exertions.

To analyze the aversive and appetitive nature of effort and reward, we evaluated the baseline portion of acceptance for each participant in choice trials across effort and reward levels:

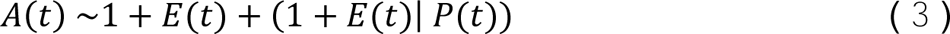

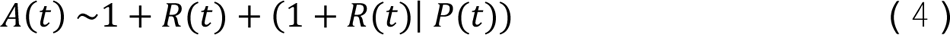

*A(t)* is the proportion of acceptance for a particular effort or reward level at the participant level *t*, *E(t)* is the effort level of the non-default option as the *n* in the *n*-back being offered, and *R(t)* is the reward corresponding to the successful completion of the non-default offer. *E(t)* and *R(t)* were both z-scored and these models were fitted via maximum likelihood estimation. A random effect of the slope was included due to the expectation that participants would have varying sensitivity to effort and reward.

### Mixed-Effects Model of Fatigue Induced Difference in Choice

To analyze the difference in behavior between the baseline and fatigue phases, we conducted a generalized linear mixed effects model:

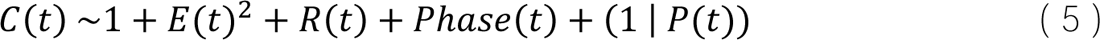

*C(t)* is the choice being made on a given trial *t* (0 = default option, 1 = non-default option), *E(t)*^2^ is the effort of the non-default option encoded as the square of the *n* in the *n*-back being offered, *R(t)* is the reward corresponding to the successful completion of the non-default effort, *Phase(t)* is a categorical variable of phase, and *P(t)* is a categorical participant identifier. Effort and reward were z-scored with their corresponding range for better direct comparison between estimated coefficients. Models were estimated with the *fitglme* function of MATLAB 2023a^33^ using the Laplace Approximation of maximum likelihood^34,35^ and an assumption of a binomial distribution of the binary response *C(t)* variable. Furthermore, the categorical variable of phase was encoded relative to the baseline phase.

### Subjective Value of Effort

Participants’ choices were fitted to structural models using maximum likelihood estimation with a sigmoid function representing the likelihood of accepting the non-default option (C = 1), where τ is the stochasticity of the participants’ choice and *SV*(*t*) is the utility function of a particular offer *t* in Eqn. 6.

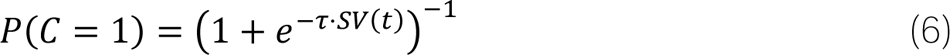

We fitted multiple models using different functions to estimate the subjective value of an offer according to the types shown in Chong. et. al.^10^.

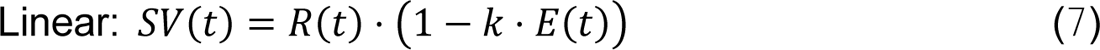

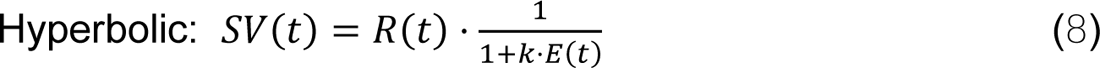

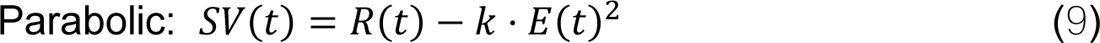

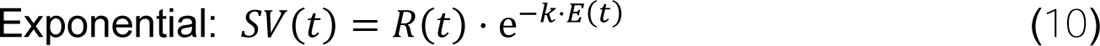

The model of best fit via both AIC and BIC was the parabolic model (Eqn. 9), where *R*(*t*) is the reward and *E*(*t*) is the effort level of the offered non-default option, while *k* is a free parameter estimated for each participant to measure their individual value of reward in the context of cognitive effort. This model of cognitive effort discounting with the *n*-back task has been validated in previous studies^36^. Models were run separately for the baseline and fatigue phases to extract *k* parameters for each participant under both conditions.

### fMRI Analysis

#### Image Preprocessing and fMRI Analysis

MRI images were preprocessed using the SPM12 software package (Wellcome Trust Centre for Neuroimaging, Institute of Neurology; London, UK)^37^. Each functional image was registered to the mean of all images, smoothed with a Gaussian kernel of 5mm FWHM and resliced, and interpolated with a 4th-degree B-Spline. Slice timing correction was performed to correct for interleaved slices every seventh slice until the end of all 48 slices, using the first slice as a reference. Images were then coregistered to the MPRAGE anatomical image using a normalized mutual information objective function with a separation of 4mm x 2mm. Images were segmented with SPM12 Tissue Probability Maps. Finally, images were normalized and smoothed with an 8mm x 8mm x 8mm FWHM Gaussian Kernel. The exact preprocessing pipeline, with all specified settings, is available in the Open Science Framework repository (https://osf.io/wfnu9/)^38^.

A general linear model (GLM) was used to estimate participant-specific (first-level), voxel-wise, statistical parametric maps (SPMs) from the fMRI data. The GLM included categorical box-car regressors beginning at the time of trial presentation and ending when a choice was indicated, for the baseline and fatigue choice phases. Each categorical regressor included orthogonalized parametric modulators of subjective chosen and unchosen value. Each participant’s subjective value was calculated by transforming effort values using their estimated parabolic effort discounting model. Trials with missing choice responses were modeled as a separate nuisance regressor. Another categorical regressor was included to model the fatiguing *n*-back blocks. The *n*-back condition was modeled as a single block with duration beginning at the time of the *n*-back onset and ending at the completion of the last *n*-back trial. This condition included a parametric modulator associated with *n*-back block number. Regressors modeling head motion as derived from the affine part of the realignment procedure were included in the model.

With these first-level models, we created group models (second-level) to test brain areas that were sensitive to subjective cognitive effort/reward value. This was done by creating contrasts with the parametric modulators for the chosen value at the time of choice. To test for regions of the brain sensitive to subjective value, irrespective of fatigue state, we created a contrast that selected the subjective chosen value parametric modulators for both the baseline and fatigue choice conditions. We also tested for regions of the brain in which the chosen subjective value was sensitive to changes in bodily state induced by fatigue by taking the difference between the chosen value in fatigue and baseline choice phases. In addition, we tested for changes in brain activity that were sensitive to repeated fatiguing cognitive exertion by selecting the parametric modulator for block number in the fatiguing *n*-back categorical regressor.

To conduct statistical inference, we extract signal change from independent regions of interest (ROI) in the brain using the MarsBaR^39^ toolbox for SPM12^37^. ROIs during choice were independently defined by using the conjunction of pre-determined functional regions (regions including the terms “vmPFC”, “dACC”, NAc”, and “aIns” in its labeling) from a task-based fMRI atlas^40^ and predictive maps from NeuroQuery^41^ associated with the search term “effort choice”. ROIs set for activity associated with increasing *n-*back were similarly determined using the intersection of regions of the a task-based fMRI atlas^40^ labeled as the “dlPFC” and the predictive map from NeuroQuery^41^ for the search term “working memory task”.

For Psychophysiological Interaction (PPI) analysis, we used a simplified GLM that unified all activity during the time of choice by removing the chosen and unchosen value parametric modulators from the originally described GLM. A physiological seed was defined by a task-based fMRI atlas^40^ that included the terms “aIns” in conjunction with “effort choice” predictive map from NeuroQuery^41^. Timeseries data was extracted using the GLMs explained above, and contrasts associated with the difference between fatigue and baseline choice BOLD signal (Fatigue Choice > Baseline Choice). The same ROI used to evaluate activity during the *n*-back task was also used here to evaluate the activity connected to the rIns during fatigue at the time of choice. The exact analysis pipeline, with all specified settings, is available in the Open Science Framework repository (https://osf.io/wfnu9/)^38^.

## Data Availability

The datasets generated and analyzed during this study are available in the Open Science Framework repository (https://osf.io/wfnu9/)^38^. Included is an imaging data set in accordance with the Brain Imaging Data Structure standard^42^ and validated with the BIDS-Validator tool^43^.

## Code Availability

The code used to collect and analyze data during this study are available in the Open Science Framework repository (https://osf.io/wfnu9/)^38^.

## Notes

https://osf.io/wfnu9/

### Competing Interest Statement

The authors have declared no competing interest.

