## Supplemental Information for "The Neurobiology of Cognitive Fatigue and Its Influence on Effort-Based Choice"

Vikram S. Chib

716 North Broadway

Rm 241

Baltimore, MD 21205, USA

443-923-2716

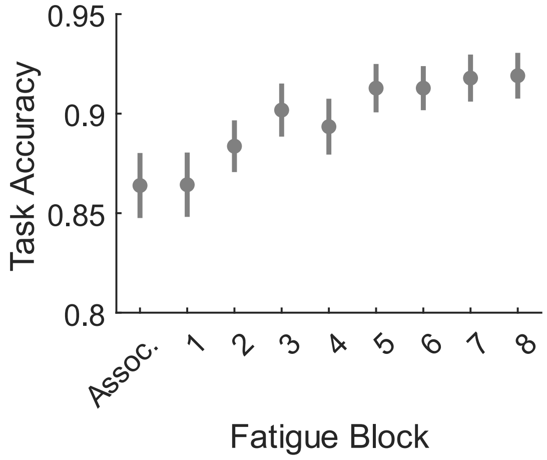

***Figure S1. 3-Back Task Performance.*** *Accuracy of identifying targets and non-target letters on the 3-Back task throughout the association and fatigue phases. Means of the accuracy are plotted with error bars representing standard error. We conducted a linear mixed-effects model to verify the effect of fatigue block on task accuracy ($\beta_{Block}$ = 0.007, SE = 0.001, t¬222 = 6.95, P << 0.001).*

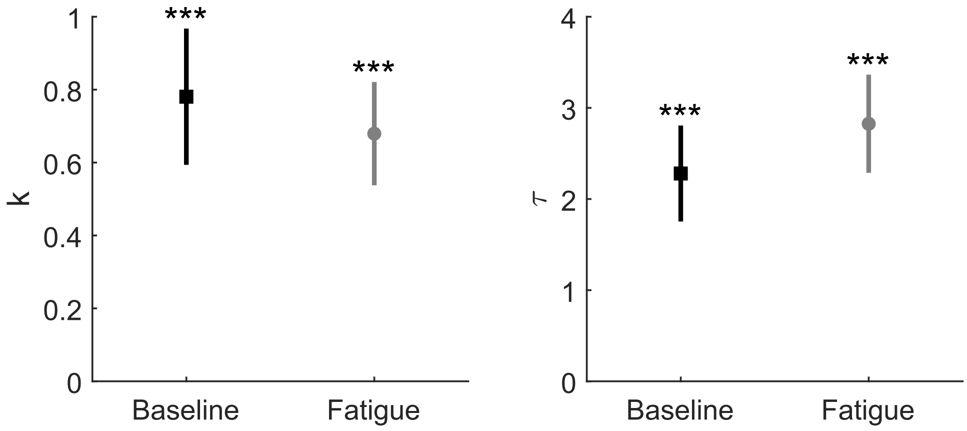

***Figure S2. Effect of Fatigue on Model Parameters.*** *We found no significant difference in the model parameter estimates between the two phases (k: $t_{27}$ = 1.094, P = 0.283; τ: $t_{27}$ = -0.934, P = 0.358).*

***Table S1. Mental Demand Affected by n-back Level.*** *Linear mixed effects model of n-back level on mental demand ratings.*

**Formula:** rating ~ 1 + lvl + (1 + lvl | subj)

**Model Information:**

| | |
|---|---|
| Number of observations | 168 |
| Fixed effects coefficients | 2 |
| Random effects coefficients | 56 |
| Covariance parameters | 4 |

**Model Fit Statistics:**

| AIC | BIC | Log Likelihood | Deviance |
|---|---|---|---|
| 374 | 393 | -181 | 362 |

**Fixed Effect Coefficients:**

| Name | Estimate | SE | tStat | DF | pValue | Lower 95% CI | Upper 95% CI |
|---|---|---|---|---|---|---|---|
| Intercept | 0.000 | 0.055 | 0.000 | 166 | 1.000 | -0.108 | 0.108 |
| lvl | 0.702 | 0.055 | 12.768 | 166 | 1.9E-26 | 0.593 | 0.810 |

***Table S2. Fatigue Ratings Increase with Additional Fatigue Blocks.*** *Linear mixed effects model of fatiguing block number on fatigue ratings.*

**Formula:** fatRating ~ 1 + block + (1 + block | subj)

**Model Information:**

| | |
|---|---|
| Number of observations | 224 |
| Fixed effects coefficients | 2 |
| Random effects coefficients | 56 |
| Covariance parameters | 4 |

**Model Fit Statistics:**

| AIC | BIC | Log Likelihood | Deviance |
|---|---|---|---|
| 327 | 348 | -158 | 315 |

**Fixed Effect Coefficients:**

| Name | Estimate | SE | tStat | DF | pValue | Lower 95% CI | Upper 95% CI |
|---|---|---|---|---|---|---|---|
| Intercept | -6.66E-15 | 1.53E-01 | -4.35E-14 | 222 | 1 | -0.302 | 0.302 |
| block | 3.98E-01 | 5.72E-02 | 6.95E+00 | 222 | 4E-11 | 0.285 | 0.510 |

***Table S3. Task Performance Increases with Fatigue Blocks.*** *Linear mixed effects model of working memory task performance with increasing fatigue blocks.*

**Formula:** AveAccuracyRate ~ 1 + BlockNum + (1 | subj))

**Model Information:**

| | |
|---|---|
| Number of observations | 224 |
| Fixed effects coefficients | 2 |
| Random effects coefficients | 28 |
| Covariance parameters | 2 |

**Model Fit Statistics:**

| AIC | BIC | Log Likelihood | Deviance |
|---|---|---|---|
| 374 | 393 | -181 | 362 |

**Fixed Effect Coefficients:**

| Name | Estimate | SE | tStat | DF | pValue | Lower 95% CI | Upper 95% CI |
|---|---|---|---|---|---|---|---|
| Intercept | 0.868 | 0.012 | 71.778 | 222 | 1.31E-155 | 8.44E-01 | 8.92E-01 |
| BlockNum | 0.007 | 0.001 | 6.955 | 222 | 3.90E-11 | 5.18E-03 | 9.27E-03 |

***Table S4. Cognitive Effort Decreases Acceptance of Non-Default Option.*** *Linear mixed effects model of cognitive effort on accepting the non-default option.*

**Formula:** AcceptanceRate ~ 1 + EffortLvl + (1 + EffortLvl | subj)

**Model Information:**

| | |
|---|---|
| Number of observations | 140 |
| Fixed effects coefficients | 2 |
| Random effects coefficients | 56 |
| Covariance parameters | 4 |

**Model Fit Statistics:**

| AIC | BIC | Log Likelihood | Deviance |
|---|---|---|---|
| 320 | 337 | -154 | 308 |

**Fixed Effect Coefficients:**

| Name | Estimate | SE | tStat | DF | pValue | Lower 95% CI | Upper 95% CI |
|---|---|---|---|---|---|---|---|
| Intercept | -4.87E-16 | 0.097 | -5.04E-15 | 138 | 1 | -0.191 | 0.191 |
| EffortLvl | -0.621 | 0.068 | -9.181 | 138 | 6E-16 | -0.754 | -0.487 |

***Table S5. Reward Increasees Acceptance of Non-Default Option.*** *Linear mixed effects model of reward on accepting the non-default option.*

**Formula:** AcceptanceRate ~ 1 + RewardLvl + (1 + RewardLvl | subj)

**Model Information:**

| | |
|---|---|
| Number of observations | 224 |
| Fixed effects coefficients | 2 |
| Random effects coefficients | 56 |
| Covariance parameters | 4 |

**Model Fit Statistics:**

| AIC | BIC | Log Likelihood | Deviance |
|---|---|---|---|
| 391 | 411 | -189 | 379 |

**Fixed Effect Coefficients:**

| Name | Estimate | SE | tStat | DF | pValue | Lower 95% CI | Upper 95% CI |
|---|---|---|---|---|---|---|---|
| Intercept | 1.34E-14 | 0.101 | 1.32E-13 | 222 | 1 | -0.200 | 0.200 |
| RewardLvl | 0.649 | 0.072 | 8.959 | 222 | 1E-16 | 0.506 | 0.792 |

***Table S6. Fewer Non-Default Options Accepted Due to the Fatigue Phase.*** *General linear mixed effects model of all components of choice and phase*

**Formula:** C ~ E_sq + R + fatPhase + (1 + E_sq + R | subj))

**Model Information:**

| | |
|---|---|
| Number of observations | 4437 |
| Fixed effects coefficients | 3 |
| Random effects coefficients | 84 |
| Covariance parameters | 6 |

**Model Fit Statistics:**

| AIC | BIC | Log Likelihood | Deviance |
|---|---|---|---|
| 2893 | 2951 | -1437 | 2875 |

**Fixed Effect Coefficients:**

| Name | Estimate | SE | tStat | DF | pValue | Lower 95% CI | Upper 95% CI |
|---|---|---|---|---|---|---|---|
| E_sq | -2.089 | 0.226 | -9.240 | 4434 | 3.72E-20 | -2.532 | -1.645 |
| R | 2.329 | 0.216 | 10.784 | 4434 | 8.74E-27 | 1.905 | 2.752 |
| fatPhase | -0.349 | 0.097 | -3.598 | 4434 | 3.24E-04 | -0.540 | -0.159 |

***Table S7. Model Fit Criteria for Subjective Value Models.***

| *Model Type* | *BIC* |
|---|---|
| *Linear:* $SV(t) = R(t) \cdot \left(1 - k \cdot E(t)\right)$ | 1166 |
| *Hyperbolic:* $SV(t) = R(t) \cdot \frac{1}{1+k \cdot E(t)}$ | 1534 |
| *Parabolic:* $SV(t) = R(t) - k \cdot E(t)^2$ | 909 |
| *Exponential:* $SV(t) = R(t) \cdot e^{-k \cdot E(t)}$ | 1387 |

***Table S8. Active Regions for Chosen Value.*** *Brain regions of clusters with a significant increase in fMRI Bold Signal associated with increases in the subjective value of the chosen option at the time of choice in both the baseline and fatigue phase (p < 0.001, k = 10, uncorrected).*

| | | Peak MNI Coordinates (mm) | | | |
|---|---|---|---|---|---|
| **Brain Region (AAL3)** | **Laterality** | **x** | **y** | **z** | **Peak *t*-value** |
| Insula | L | -32 | 24 | 6 | 6.43 |
| Mediodorsal medial magnocellular | L | -6 | -16 | 2 | 5.84 |
| Mediodorsal medial magnocellular | R | 2 | -12 | 4 | 5.27 |
| Inferior occipital gyrus | L | -42 | -66 | -6 | 6.07 |
| Middle occipital gyrus | L | -22 | -92 | 10 | 5.90 |
| Fusiform gyrus | L | -46 | -54 | -18 | 5.31 |
| Middle temporal gyrus | R | 50 | -36 | -14 | 5.96 |
| Fusiform gyrus | R | 36 | -50 | -20 | 5.41 |
| Fusiform gyrus | R | 36 | -58 | -14 | 5.19 |
| Inferior occipital gyrus | R | 32 | -82 | -14 | 5.01 |
| Supplementary motor area | L | -10 | -2 | 54 | 5.28 |
| Precentral gyrus | L | -20 | -16 | 54 | 4.37 |
| Middle cingulate & paracingulate gyri | L | -14 | -14 | 44 | 4.35 |
| Inferior frontal gyrus, triangular part | R | 36 | 26 | 10 | 5.18 |
| Anterior cingulate cortex, supracallosal | R | 24 | 30 | 16 | 4.39 |
| Inferior frontal gyrus, triangular part | R | 42 | 20 | 14 | 3.51 |
| Anterior cingulate cortex, pregenual | R | 10 | 40 | 16 | 5.09 |
| Anterior cingulate cortex, pregenual | L | -4 | 42 | 14 | 4.13 |
| IFG pars orbitalis, | R | 36 | 28 | -12 | 5.08 |
| Insula | R | 36 | 16 | -14 | 4.91 |
| IFG pars orbitalis, | R | 42 | 36 | -6 | 4.15 |
| Anterior cingulate cortex, pregenual | L | -8 | 54 | 2 | 4.81 |
| Anterior cingulate cortex, pregenual | L | -8 | 40 | -2 | 4.76 |
| Anterior cingulate cortex, supracallosal | L | -2 | 30 | 8 | 3.96 |
| Anterior cingulate cortex, supracallosal | R | 8 | 18 | 26 | 4.63 |
| Postcentral gyrus | L | -36 | -26 | 48 | 4.38 |
| Postcentral gyrus | L | -46 | -26 | 50 | 3.68 |
| Lenticular nucleus, Putamen | R | 30 | -2 | 6 | 4.32 |
| Lenticular nucleus, Pallidum | R | 26 | -12 | -2 | 4.10 |
| Precentral gyrus | R | 58 | 6 | 38 | 4.19 |
| Middle occipital gyrus | R | 32 | -68 | 26 | 4.18 |
| Superior occipital gyrus | R | 26 | -62 | 32 | 3.73 |
| Precentral gyrus | L | -36 | -6 | 46 | 4.18 |
| Precentral gyrus | L | -50 | 0 | 42 | 4.17 |
| Precentral gyrus | L | -48 | -4 | 54 | 3.79 |
| Middle temporal gyrus | L | -58 | -38 | 4 | 4.06 |
| Middle occipital gyrus | L | -30 | -82 | 20 | 4.06 |
| Middle occipital gyrus | L | -30 | -74 | 28 | 3.54 |
| Inferior parietal gyrus | R | 32 | -52 | 48 | 4.04 |

| *(Continued)* | | Peak MNI Coordinates (mm) | | | |
|---|---|---|---|---|---|
| **Brain Region (AAL3)** | **Laterality** | **x** | **y** | **z** | **Peak *t*-value** |
| Inferior parietal gyrus | L | -32 | -52 | 40 | 4.03 |
| Postcentral gyrus | L | -20 | -44 | 56 | 4.01 |
| Postcentral gyrus | L | -20 | -36 | 58 | 3.65 |
| Middle temporal gyrus | R | 50 | -42 | 6 | 4.02 |
| Angular gyrus | L | -48 | -52 | 26 | 3.95 |
| Supramarginal gyrus | L | -48 | -46 | 32 | 3.86 |
| Inferior parietal gyrus | L | -44 | -42 | 40 | 3.79 |
| Precentral gyrus | L | -30 | -14 | 72 | 3.89 |
| Superior frontal gyrus, medial | R | 10 | 42 | 32 | 3.86 |
| Supramarginal gyrus | L | -44 | -40 | 26 | 3.85 |
| Superior temporal gyrus | L | -46 | -46 | 18 | 3.53 |
| Inferior frontal gyrus, opercular part | R | 52 | 14 | 16 | 3.85 |
| Inferior frontal gyrus, opercular part | R | 54 | 6 | 20 | 3.84 |
| Rolandic operculum | R | 52 | 4 | 12 | 3.79 |
| Paracentral lobule | L | -12 | -30 | 52 | 3.82 |
| Pulvinar lateral | L | -26 | -28 | 8 | 3.80 |
| Crus II of cerebellar hemisphere | L | -10 | -80 | -34 | 3.80 |
| Inferior frontal gyrus, opercular part | R | 38 | 14 | 34 | 3.70 |
| Middle temporal gyrus | L | -54 | -24 | -4 | 3.68 |
| Middle cingulate & paracingulate gyri | L | -6 | -34 | 34 | 3.68 |
| Crus II of cerebellar hemisphere | R | 16 | -74 | -24 | 3.68 |
| Supramarginal gyrus | L | -56 | -30 | 34 | 3.67 |
| Cuneus | R | 14 | -72 | 38 | 3.61 |
| Superior occipital gyrus | R | 22 | -74 | 40 | 3.60 |

***Table S9. Active Regions for Difference in Chosen Value between Fatigue and Baseline.*** *Brain regions with a significant increase in fMRI Bold Signal associated with difference in the subjective value of the chosen option at the time of choice in the between the fatigue and baseline phase ($p < 0.005$, $k = 10$, uncorrected).*

| | | Peak MNI Coordinates (mm) | | | |
|---|---|---|---|---|---|
| **Brain Region (AAL3)** | **Laterality** | **x** | **y** | **z** | **Peak *t*-value** |
| Caudate nucleus | R | 16 | -2 | 24 | 4.16 |
| Caudate nucleus | R | 16 | -12 | 26 | 3.66 |
| Anterior cingulate cortex, pregenual | R | 10 | 38 | 26 | 4.12 |
| Crus II of cerebellar hemisphere | L | -8 | -26 | -34 | 4.03 |
| Lenticular nucleus, Putamen | L/R | 0 | -24 | -34 | 3.96 |
| Anterior cingulate cortex, pregenual | L | -6 | 40 | 22 | 4.03 |
| Superior frontal gyrus, dorsolateral | L | -24 | 60 | 8 | 3.99 |
| Superior frontal gyrus, dorsolateral | L | -18 | 56 | 14 | 3.53 |
| Insula | R | 42 | 16 | -8 | 3.60 |
| Insula | R | 34 | 16 | -12 | 3.56 |

***Table S10. Active Regions Associated with Increasing N-Back Exertion.*** *Brain regions with a significant increase in fMRI Bold Signal associated with the increase of N-Back tasks experienced during the time N-Back task performance (p < 0.001, uncorrected).*

| | | Peak MNI Coordinates (mm) | | | |
|---|---|---|---|---|---|
| **Brain Region (AAL3)** | **Laterality** | **x** | **y** | **z** | **Peak *t*-value** |
| Caudate nucleus | R | 18 | 16 | 6 | 6.09 |
| Lenticular nucleus, Pallidum | R | 18 | 0 | -2 | 5.55 |
| Lenticular nucleus, Putamen | R | 24 | 14 | -2 | 4.30 |
| Middle frontal gyrus | R | 40 | 18 | 52 | 5.56 |
| Inferior frontal gyrus, triangular part | R | 42 | 20 | 28 | 5.06 |
| Middle frontal gyrus | R | 32 | 22 | 56 | 4.58 |
| Precentral gyrus | L | -38 | 4 | 32 | 5.35 |
| Inferior frontal gyrus, triangular part | L | -50 | 22 | 26 | 5.06 |
| Precentral gyrus | L | -42 | 8 | 48 | 4.57 |
| Superior frontal gyrus, dorsolateral | L | -14 | 32 | 54 | 5.02 |
| Superior frontal gyrus, medial | L | -8 | 54 | 36 | 4.11 |
| Superior frontal gyrus, dorsolateral | L | -14 | 50 | 40 | 3.99 |
| Superior temporal gyrus | R | 62 | -8 | -8 | 4.91 |
| Paracentral lobule | L | -2 | -24 | 66 | 4.88 |
| Paracentral lobule | L | -12 | -38 | 74 | 4.51 |
| Paracentral lobule | L | -4 | -32 | 72 | 4.28 |
| Crus II of cerebellar hemisphere | R | 34 | -62 | -38 | 4.69 |
| Crus II of cerebellar hemisphere | R | 42 | -72 | -34 | 4.26 |
| Red nucleus | R | 4 | -20 | -8 | 4.59 |
| Posterior orbital gyrus | R | 44 | 30 | -14 | 4.55 |
| IFG pars orbitalis, | R | 46 | 40 | -10 | 3.72 |
| Posterior orbital gyrus | R | 36 | 24 | -20 | 3.45 |
| Inferior temporal gyrus | L | -44 | 4 | -36 | 4.45 |
| Inferior frontal gyrus, triangular part | L | -44 | 36 | 0 | 4.35 |
| Inferior frontal gyrus, triangular part | L | -48 | 32 | 6 | 4.26 |
| IFG pars orbitalis, | L | -42 | 40 | -10 | 3.93 |
| Inferior frontal gyrus, triangular part | R | 52 | 40 | 8 | 4.25 |
| Inferior frontal gyrus, triangular part | R | 54 | 34 | 14 | 3.91 |
| Inferior frontal gyrus, triangular part | R | 44 | 32 | 8 | 3.54 |
| Inferior parietal gyrus | L | -46 | -52 | 52 | 4.25 |
| Inferior parietal gyrus | L | -48 | -46 | 44 | 3.88 |
| Inferior parietal gyrus | L | -38 | -60 | 52 | 3.66 |
| Inferior occipital gyrus | L | -20 | -90 | -6 | 4.22 |
| Insula | R | 34 | -6 | 24 | 4.20 |
| Superior frontal gyrus, dorsolateral | R | 26 | 46 | -6 | 4.15 |
| Superior frontal gyrus, medial orbital | R | 18 | 44 | -6 | 4.06 |
| Insula | R | 28 | -26 | 14 | 4.12 |

| *(Continued)* | | Peak MNI Coordinates (mm) | | | |
|---|---|---|---|---|---|
| **Brain Region (AAL3)** | **Laterality** | **x** | **y** | **z** | **Peak *t*-value** |
| Inferior parietal gyrus | R | 52 | -50 | 46 | 4.09 |
| Inferior parietal gyrus | R | 56 | -58 | 40 | 4.01 |
| Postcentral gyrus | R | 18 | -36 | 58 | 4.05 |
| Paracentral lobule | R | 14 | -26 | 58 | 3.92 |
| Temporal pole: superior temporal gyrus | R | 46 | 12 | -24 | 4.02 |
| Temporal pole: middle temporal gyrus | R | 38 | 8 | -36 | 4.00 |
| Superior frontal gyrus, medial | L | 0 | 56 | 18 | 3.98 |
| Superior frontal gyrus, dorsolateral | R | 14 | 42 | 48 | 3.97 |
| Superior frontal gyrus, dorsolateral | R | 18 | 48 | 44 | 3.52 |
| Superior frontal gyrus, medial | L | -16 | 58 | 0 | 3.94 |
| Fusiform gyrus | L | -42 | -70 | -18 | 3.89 |
| Ventral striatum | L | -4 | 12 | -4 | 3.88 |
| Middle temporal gyrus | R | 66 | -42 | -4 | 3.87 |
| Posterior orbital gyrus | L | -44 | 26 | -14 | 3.83 |
| Middle temporal gyrus | L | -52 | 4 | -26 | 3.82 |
| Middle occipital gyrus | L | -36 | -68 | 36 | 3.80 |
| Crus II of cerebellar hemisphere | R | 12 | -42 | -28 | 3.77 |
| Calcarine fissure and surrounding cortex | R | 26 | -94 | 0 | 3.74 |
| Crus II of cerebellar hemisphere | R | 20 | -84 | -38 | 3.72 |
| Crus II of cerebellar hemisphere | R | 12 | -76 | -38 | 3.52 |
| Middle temporal gyrus | L | -54 | -32 | -8 | 3.58 |
| Angular gyrus | L | -52 | -60 | 26 | 3.51 |

***Table S11. Active Regions in PPI Analysis.*** *Brain regions indicating a connection with the dlPFC working memory activity at the time of choice in the fatigue phase (p < 0.001, uncorrected).*

| | | Peak MNI Coordinates (mm) | | | |
|---|---|---|---|---|---|
| **Brain Region (AAL3)** | **Laterality** | **x** | **y** | **z** | **Peak *t*-value** |
| Precentral gyrus | L | -28 | -24 | 48 | 6.92 |
| Precentral gyrus | L | -32 | -14 | 48 | 6.48 |
| Supplementary motor area | R | 8 | -10 | 66 | 6.23 |
| Lingual gyrus | R | 14 | -72 | 2 | 6.23 |
| Crus II of cerebellar hemisphere | L | -22 | -62 | -58 | 5.80 |
| Inferior temporal gyrus | L | -50 | -62 | -8 | 5.67 |
| Middle temporal gyrus | R | 58 | -36 | -8 | 5.22 |
| Middle temporal gyrus | R | 56 | -44 | 0 | 3.69 |
| Ventral striatum | R | 12 | 10 | -6 | 4.66 |
| Superior frontal gyrus, dorsolateral | L | -24 | 42 | -6 | 4.64 |
| Middle frontal gyrus | L | -32 | 54 | 6 | 4.24 |
| Fusiform gyrus | L | -34 | -26 | -20 | 4.54 |
| Caudate nucleus | L | -14 | 4 | 16 | 4.41 |
| Anterior cingulate cortex, pregenual | L | -6 | 38 | 24 | 4.41 |
| Superior frontal gyrus, medial | R | 6 | 48 | 28 | 4.38 |
| Anterior cingulate cortex, supracallosal | L | -14 | 26 | 24 | 4.36 |
| IFG pars orbitalis, | L | -34 | 32 | -8 | 4.31 |
| Ventral striatum | L | -8 | 18 | -8 | 4.26 |
| Lenticular nucleus, Putamen | L | -18 | 16 | -4 | 3.79 |
| Superior frontal gyrus, medial orbital | R | 6 | 60 | -2 | 4.23 |
| Superior frontal gyrus, medial | R | 2 | 64 | 12 | 4.16 |
| Superior frontal gyrus, medial | L | 0 | 64 | 22 | 4.16 |
| Crus II of cerebellar hemisphere | R | 24 | -64 | -56 | 4.19 |
| Middle temporal gyrus | R | 40 | -68 | 8 | 4.16 |
| Middle temporal gyrus | R | 46 | -56 | 10 | 4.13 |
| Middle temporal gyrus | R | 52 | -50 | 12 | 3.58 |
| Olfactory cortex | L | -14 | 4 | -14 | 4.12 |
| Caudate nucleus | L | -18 | 18 | 10 | 4.11 |
| Superior frontal gyrus, medial | R | 4 | 60 | 34 | 4.05 |
| Crus II of cerebellar hemisphere | R | 22 | -48 | -52 | 4.01 |
| Fusiform gyrus | R | 46 | -50 | -20 | 4.00 |
| Fusiform gyrus | R | 44 | -42 | -22 | 3.46 |
| Inferior frontal gyrus, triangular part | L | -46 | 32 | 16 | 3.98 |
| Inferior frontal gyrus, triangular part | L | -38 | 38 | 14 | 3.80 |
| Superior parietal gyrus | L | -14 | -72 | 44 | 3.98 |
| Middle frontal gyrus | L | -24 | 46 | 32 | 3.96 |
| Anterior cingulate cortex, supracallosal | R | 14 | 32 | 24 | 3.93 |
| Anterior cingulate cortex, supracallosal | R | 10 | 36 | 14 | 3.68 |
| Parahippocampal gyrus | R | 30 | -20 | -20 | 3.90 |

| *(Continued)* | | Peak MNI Coordinates (mm) | | | |
| --- | --- | --- | --- | --- | --- |
| **Brain Region (AAL3)** | **Laterality** | **x** | **y** | **z** | **Peak *t*-value** |
| Inferior frontal gyrus, triangular part | L | -44 | 30 | 28 | 3.87 |
| Middle temporal gyrus | R | 58 | -64 | 12 | 3.79 |
| Caudate nucleus | R | 18 | 20 | 4 | 3.75 |
| Middle temporal gyrus | R | 52 | -62 | -2 | 3.67 |
| Middle temporal gyrus | R | 52 | -70 | 2 | 3.52 |
| SupraMarginal gyrus | R | 66 | -24 | 28 | 3.62 |
